# Long-term effects of moderate concussive brain injury during adolescence on synaptic and tonic GABA currents in dentate projection neurons

**DOI:** 10.1101/2021.08.15.456415

**Authors:** Akshay Gupta, Archana Proddutur, Fatima S. Elgammal, Vijayalakshmi Santhakumar

**Author notes:** **Author Contact:** Akshay Gupta, Archana Proddutur, Fatima Elgammal, Vijayalakshmi Santhakumar. Correspondence: **Vijayalakshmi Santhakumar, PhD**, Department of Molecular Cell and Systems Neuroscience, University of California, Riverside, 3401 Watkins Drive (Rm 1308 Spieth Hall) Riverside, CA 92521.

## Abstract

Progressive physiological changes in the hippocampal dentate gyrus circuits following traumatic brain injury contribute to temporal evolution of neurological sequelae. Although early posttraumatic changes in dentate synaptic and extrasynaptic GABA currents have been reported, whether they evolve over time and remain distinct between the two projection neuron classes, granule cells and semilunar granule cells, has not been evaluated. We examined changes in tonic GABA currents and spontaneous inhibitory postsynaptic currents (sIPSCs) and in dentate projection neurons one and three month after moderate concussive fluid percussion injury (FPI) in adolescent rats. Granule cell tonic GABA current amplitude remained elevated up to one month after FPI, but decreased to levels comparable to age-matched controls by three months postinjury. Granule cell sIPSC frequency, which we previously reported to be increased one week after FPI, remained higher than in age-matched controls at one month and was significantly reduced three months after FPI. In contrast to the early decrease, tonic GABA current amplitude and sIPSC frequency in semilunar granule cell was not different from controls three months after FPI. The switch in granule cell inhibitory inputs from early increase to subsequent decrease could contribute to the delayed emergence of cognitive deficits and seizure susceptibility after brain injury.

## Introduction

Traumatic brain injury (TBI) leads to diverse consequences including impaired memory and reasoning, depression, anxiety as well as enhanced risk for epilepsies and Alzheimer’s Disease (LoBue et al., 2019; Faden et al., 2021). TBI is a lifelong disease with adverse outcomes that evolve over months to years, highlighting the need to understand progressive changes in cellular and circuit function following brain injury (Marshall et al., 2015). The hippocampal dentate gyrus is a focus of cellular pathology and functional changes following brain injury in humans and various experimental models of TBI (Kharatishvili et al., 2006; Hunt et al., 2010; Villasana et al., 2015; Meier et al., 2016; Neuberger et al., 2017a; Neuberger et al., 2017b; Wolf et al., 2017; Parga Becerra et al., 2021). The dentate gyrus is a crucial gateway to the hippocampal circuit, serving as a locus for memory processing and as a check against excessive excitability and reentrant epileptiform activity (Dengler and Coulter, 2016). Strong synaptic and extrasynaptic inhibition of dentate projection neurons contributes to their sparse activity and is known to be disrupted in TBI and epilepsies (Peng et al., 2004; Rajasekaran et al., 2010; Pavlov et al., 2011; Gupta et al., 2012; Boychuk et al., 2016; Kahn et al., 2019; Parga Becerra et al., 2021). Studies examining injury-induced changes in inhibition in dentate granule cells (GCs), the major projection neuronal subtype, days to weeks after trauma have identified changes in synaptic and extrasynaptic GABA_A_ currents which differ between experimental injury models and on the basis of injury severity (Santhakumar et al., 2001; Pavlov et al., 2011; Gupta et al., 2012; Boychuk et al., 2016). While long-term (> 4 week) posttraumatic changes in GC inhibition have been examined, the results are varied, with persistent decreases in both tonic and synaptic GABA_A_ currents after cortical impact injury (CCI) (Boychuk et al., 2016; Parga Becerra et al., 2021) and reduced synaptic inhibition while tonic GABA_A_ currents remained unchanged after severe concussion (Pavlov et al., 2011). We previously reported an increase in GC synaptic and tonic GABA currents one week after moderate concussive brain injury in adolescent rats (Gupta et al., 2012). This clinically-relevant adolescent concussive injury paradigm impairs working memory performance one to four weeks post-injury followed by an apparent recovery yet heightened risk for seizures one to three months after injury (Neuberger et al., 2017b; Korgaonkar et al., 2020b; Korgaonkar et al., 2020a). Since GABAergic signaling which critically regulates dentate memory function and epileptogenesis (Dengler and Coulter, 2016) undergoes developmental plasticity spanning this period (Kapur and Macdonald, 1999; Gupta et al., 2020), it is important to determine if GC inhibition undergoes progressive changes after concussive TBI.

In addition to GCs, we reported that brain injury impacts inhibition in semilunar granule cells (SGCs), a sparse morphologically distinct dentate projection neuron (Williams et al., 2007; Gupta et al., 2012; Gupta et al., 2020). Interestingly, both tonic and synaptic GABA currents in SGCs are decreased one week after FPI, which contrasts with increases observed in GCs at the same time (Gupta et al., 2012). Crucially, SGCs have been proposed to support feedback inhibition of GCs needed to maintain sparse activity and contribute to cellular memory representations (Larimer and Strowbridge, 2010; Erwin et al., 2020). Since changes in tonic GABA currents enhance SGC excitability early after brain injury (Gupta et al., 2012), changes in SGC inhibition could contribute to posttraumatic memory dysfunction. While SGC tonic GABA currents are greater than in GCs during adolescence, they undergo a developmental decline into adulthood (Gupta et al., 2012; Gupta et al., 2020). Thus, whether the early decrease in SGC tonic GABA currents after injury persists at later time points when SGC tonic GABA currents have declined remains to established. Since GCs and SGCs show opposing changes in inhibition early after FPI (Gupta et al., 2012) and are proposed to play distinct roles in dentate feedback inhibition and memory processing (Larimer and Strowbridge, 2010; Erwin et al., 2020), it is crucial to understand how injury-induced changes in inhibition evolve in these dentate projection neuron subtypes. This study examined the long-term changes in tonic and synaptic GABA currents in GCs and SGCs after moderate concussive FPI in adolescent rats to determine if early cell-specific posttraumatic changes in inhibition, observed one week after moderate FPI, are maintained at later time points.

## Materials and Methods

### Animals

All experiments under IACUC protocols approved by Rutgers-NJMS and the University of California at Riverside and conformed with the ARRIVE guidelines. Wistar rats (Charles River) aged 60-70 days or 120-180 days, which were one and three months after sham or brain injury, respectively, were used in the study. Due to the potential effects of hormonal variation on GABA currents (Maguire and Mody, 2009), recordings were restricted to male rats.

### Surgery and FPI

Lateral fluid percussion injury (FPI) was performed on adolescent (postnatal day 24-26) male Wistar rats as described previously (Dixon et al., 1987; Li et al., 2015). Briefly, under ketamine-xylazine anesthesia, a 2 mm hole was trephined on the left side of the skull (3 mm posterior to bregma and 3.5 mm from lateral to sagittal suture) to expose the dura and a syringe hub with a 2.6 mm inner diameter and two anchor screws were bonded to the skull with cyanoacrylate adhesive. One day later, animals were anesthetized with isoflurane and attached to a fluid percussion device (Virginia Commonwealth University, Richmond, VA). A pendulum was dropped to deliver a brief (20 ms) 2.0 – 2.2 atm impact on the intact dura resulting in a moderate injury with reproducible pattern of hilar cell loss (Gupta et al., 2012; Li et al., 2015). Sham injured animals received identical surgery and treatment, but the pendulum was not dropped.

### Slice Physiology

The rats were anesthetized with isoflurane and decapitated. Horizontal brain slices (300μm) were prepared in ice-cold sucrose artificial-cerebrospinal fluid (sucrose-aCSF) containing the following (in mM): 85 NaCl, 75 sucrose, 24 NaHCO3, 25 glucose, 4 MgCl2, 2.5 KCl, 1.25 NaH2PO4, and 0.5 CaCl, bisected and incubated at 32 °C for 30 min in a holding chamber containing an equal volume of sucrose-aCSF and recording aCSF and subsequently were held at room temperature. The recording aCSF contained the following (in mM): 126 NaCl, 2.5 KCl, 2 CaCl2, 2 MgCl2, 1.25 NaH2PO4, 26 NaHCO3, and 10 D-glucose saturated with 95% O2 and 5% CO2 (pH 7.4). Whole-cell voltage-clamp recordings were performed at 33°C under IR-DIC visualization using MultiClamp 700B (Molecular Devices) as detailed previously (Gupta et al., 2012; Yu et al., 2016). Data were low pass filtered at 3 kHz, digitized using DigiData 1440A, and acquired at 10 kHz sampling frequency using pClamp10. Currents were recorded without added GABA or GABA transporter antagonists (Gupta et al., 2012; Yu et al., 2013). GABA currents were recorded using microelectrodes (5–7 MΩ) containing (in mM): 125 CsCl, 5 NaCl, 10 HEPES, 2 MgCl2, 0.1 EGTA, 2 Na-ATP, 0.5 Na-GTP, and 0.2% biocytin, titrated to a pH 7.25. Kynurenic acid (3 mM KynA), a glutamate receptor antagonist, was used to isolate GABA currents in cells held at −70 mV. SGCs were distinguished by large dendritic angle, multiple primary dendrites, presence of dendritic spines, and axon projecting to the hilus (Gupta et al., 2020). Recordings were discontinued if series resistance increased by >20%. Access resistance was not different between cell-types or experimental groups. Baseline recordings (in KynA) were obtained for over five minutes before perfusing GABA_A_ receptor (GABA_A_R) antagonist, bicuculline methiodide (BMI, 100μM) or gabazine (SR95531, 10 μM). Cesium-based internal solution was used to block K^+^ conductances underlying postsynaptic GABA_B_ currents. Custom macros in IgorPro7.0 were used to measure tonic GABA currents as the difference in baseline currents in blockers and for detecting sIPSCs (Gupta et al., 2012; Yu et al., 2013). All salts were purchased from Sigma-Aldrich.

### Cell morphology

Recorded slices were fixed in 4% paraformaldehyde and processed for biocytin staining using Alexa Fluor 594-conjugated streptavidin (Gupta et al., 2012; Swietek et al., 2016). Sections were imaged using a Zeiss LSM 510 confocal microscope for classification. A subset of cells were reconstructed using Neurolucida 360 (Gupta et al., 2020).

### Statistics

Cumulative probability plots of sIPSC parameters were constructed using IgorPro by pooling an equal number of sIPSCs from each cell. Wilcoxon Rank Test was conducted on data that failed tests for normalcy or equal variance. Cohen’s D test was used to estimate effect size. Student’s T-test (SigmaPlot 12.3) was used to test for statistical differences in tonic GABA currents. The summary values of physiological parameters and statistical comparisons are included in Tables 1 and 2. Sample sizes were not predetermined and conformed with those employed in the field (Table 1). Data that fell over three standard deviations outside the mean were considered outliers and rejected. Significance was set to p<0.05. Data are shown as mean±SEM (standard error of the mean) or median and interquartile range (IQR), where appropriate, and presented in Table 1.

## Results

### Post-traumatic increase in granule cell tonic GABAergic currents declines with time

Tonic GABA currents mediated by extrasynaptic receptors contribute to reduced GC excitability (Stell et al., 2003; Brickley and Mody, 2012). Granule cell tonic GABA currents are altered in experimental brain injury (Gupta et al., 2012; Boychuk et al., 2016; Parga Becerra et al., 2021). We previously reported a potentially beneficial increase in GC tonic GABA currents one week after moderate FPI 25-26 day old rats (Gupta et al., 2012), a period that parallels human adolescence in terms of neurological developmental (Semple et al., 2013; Sengupta, 2013). However, studies 1-5 months after severe FPI in adult rats did not find changes in tonic GABA currents (Pavlov et al., 2011). To determine whether developmental progression of pathology rather than model-specific differences accounted for these findings, we examined rats injured during adolescence at one and 3-4 months post-FPI. Since injury-induced changes in tonic GABA currents differ between GC and SGCs (Gupta et al., 2012), presence of 1-2 primary dendrites and compact dendritic arbors of biocytin filled cells was used to identify GCs (Fig. 1 A-B, Gupta et al., 2020). Tonic GABA current amplitude in GCs remained larger than in age-matched sham controls one-month post-FPI (Fig 1C-D and Table 1 & 2) as reported at one week (Fig. 1G, data one week post-FPI was previously reported in Gupta et al., 2012, shown shaded). However, tonic GABAergic current amplitude in GCs from rats 3-5 months after FPI was not different from age-matched controls (Fig 1E-G and Table 1 & 2). Notably, while tonic GABA currents in GCs from sham rats remained relatively stable over the period examined (1-6 months of age), there was a steady decline in tonic GABA currents from one week to three months post-FPI (Fig. 1G). These data demonstrate that the early increase in GC tonic GABA currents progressively declines to sham control levels by 3 months.

**Figure 1:**
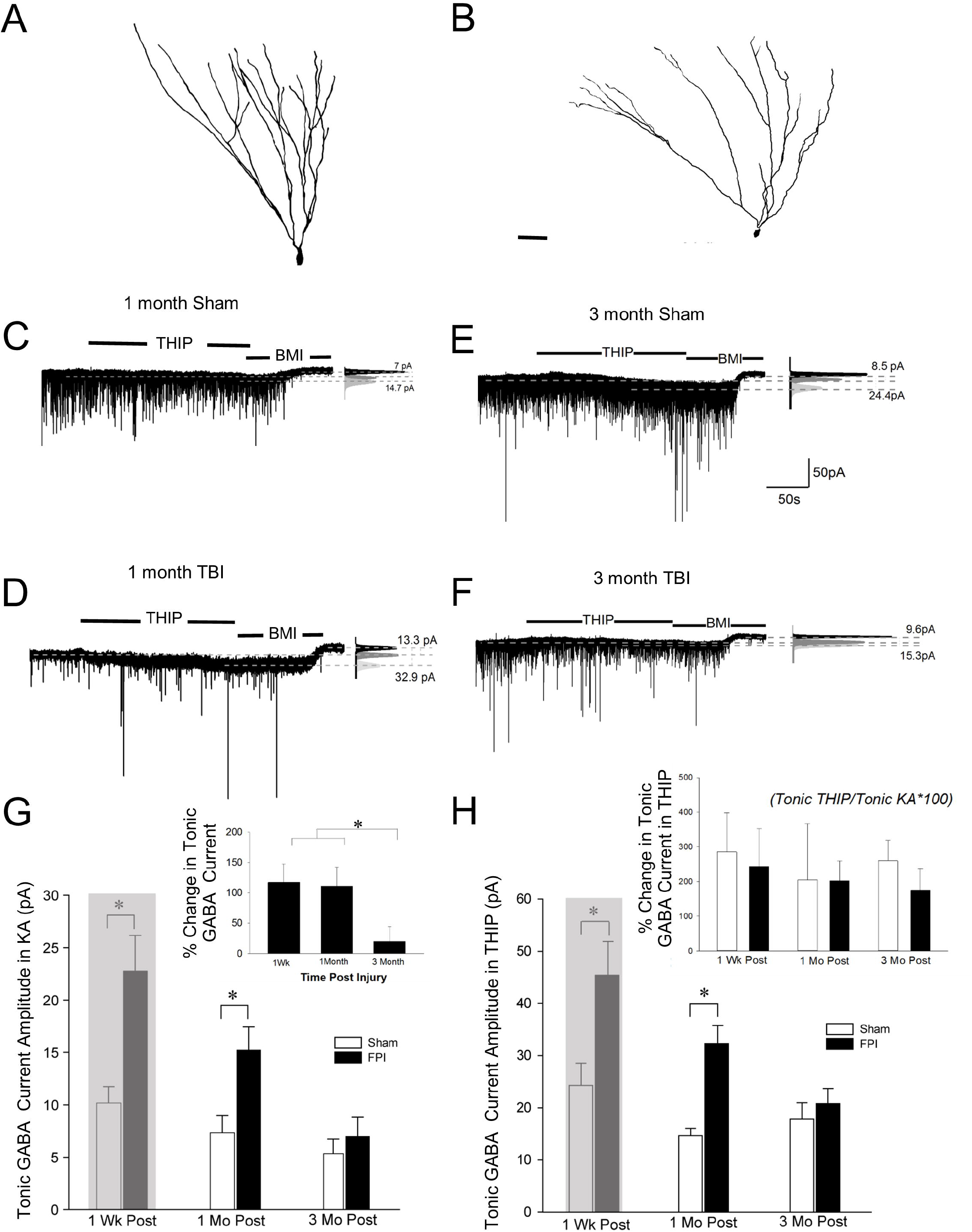
Granule cell tonic GABA current amplitude progressively declines 1 month to 3 months after brain injury. A-B. Representative Neurolucida reconstructions of GCs from rats 3 months after sham (A) and FPI (B). C-D. Representative traces from granule cell one month after sham (C) or FPI (D) illustrate tonic GABA current as the baseline current blocked by bicuculline. E-F. Representative tonic GABA current traces in GCs 3 months after sham (E) and FPI (F). Panels to the right show Gaussian fits to the positive half of histograms derived from 30 s recording periods in control conditions, in the presence of THIP (1 μM) and during the perfusion of BMI used to determine tonic current. The dashed lines indicate the Gaussian means, and the difference current is noted. G. Summary plots of tonic GABA current amplitude (pA) in granule cells from rats 1 week (data from Gupta et al., 2012 shown in grey box), 1month and 3 months after FPI. Inset: Summary of percentage change in tonic GABA current normalized to sham. H. Summary plots of tonic GABA current amplitude in granule cells measured in the presence of THIP (1 μM) from rats 1 week (data from Gupta et al., 2012 shown in gray box), 1 month and 3 months after FPI. Inset: Summary of percentage THIP induced change in tonic GABA current.

Since GABA_A_R δ subunit containing receptors mediate GC tonic GABA currents (Stell et al., 2003), we examined whether decline in GABA_A_R δ subunits contribute to progressive reduction in tonic GABA currents using THIP, a GABA_A_R agonist selective for δ subunit containing receptors. GC tonic GABA current amplitude in THIP (1μM) was higher one week (data reported in Gupta et al., 2012) and one month after FPI but were not different from sham controls by 3 months post-FPI (Figure 1H and Tables 1 and 2). However, the THIP mediated increase in tonic GABA currents, measured as the ratio of tonic GABA current amplitude in the presence and absence of THIP, was not different between experimental groups (Figure 1H inset). These data indicate that the proportion of δ containing GABA_A_R subunits is not altered by injury or time after injury and suggest that increase in GABA_A_R δ subunit contributes to elevated GC tonic GABA currents one week to one month after brain injury.

### Long-term reduction in inhibitory synaptic drive to granule cells after brain injury

We previously reported an increase in the frequency of spontaneous inhibitory postsynaptic currents (sIPSCs) in GCs one week after FPI, which remained elevated for up to five months despite an age-related decline in GC sIPSC frequency (Santhakumar et al., 2001; Gupta et al., 2012). While early increase in sIPSC was present regardless of glutamate receptor antagonism (Gupta et al., 2012), glutamatergic drive was found to contribute to long-term increase in sIPSCs after FPI (Santhakumar et al., 2001). We examined the effect of trauma on the inhibitory circuit by blocking glutamate receptors. As reported one week after FPI (Gupta et al., 2012), GC sIPSC frequency and amplitude remained elevated one month after FPI (Fig. 2A-C, Table 2). However, sIPSC 10-90% rise time and amplitude weighted decay time constant τ_decay_) were not different from age-matched controls (10-90 rise% time in ms, sham: 0.2±0.01, FPI: 0.2±0.03, p>0.05 by t-test; τ_decay_, sham: 3.4±0.1, FPI: 3.0±0.3, p>0.05 by t-test in n= 9 sham and 6 FPI). In contrast, GC sIPSC frequency was reduced 3-5 months after FPI while sIPSC amplitude was not different between groups (Fig. 2C-D, Table 2). Both 10-90% rise time and τ_decay_ of GC sIPSCs were not different between groups 3-5 months after FPI (10-90% rise time in ms, sham: 0.2±0.01, FPI: 0.2±0.013, p>0.05 by t-test; τ_decay_, sham: 2.9±0.1, FPI: 3.2±0.2, p>0.05 by t-test in n= 14 sham and 10 FPI). These data indicate an early increase followed by a progressive decline in basal interneuronal activity 3-5 months after FPI.

**Figure 2:**
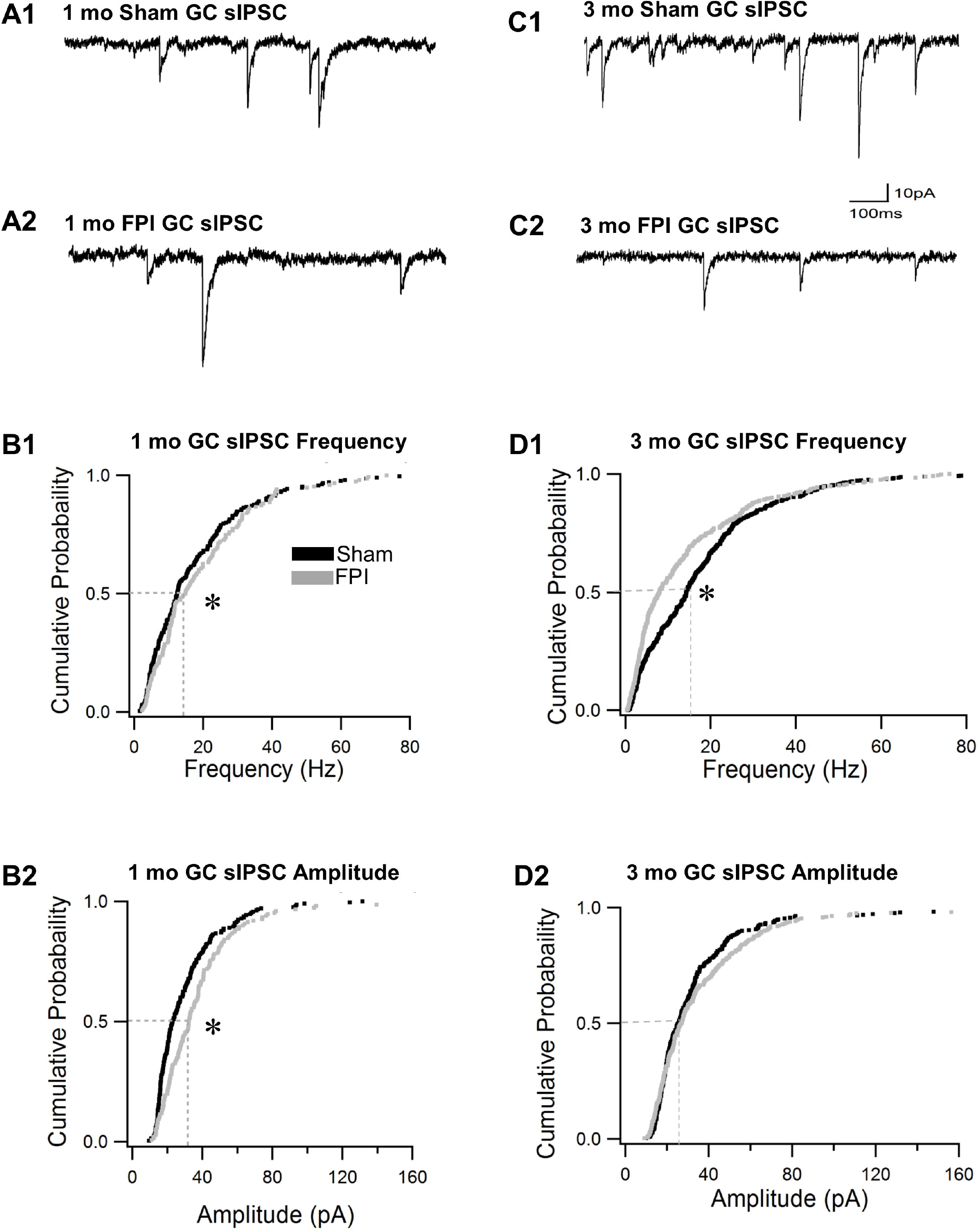
Granule cell sIPSC frequency declines over time after brain injury. A. Representative current traces illustrate sIPSCs in GCs from rats one month after sham (A1) and FPI (A2). B. Cumulative probability plots of sIPSC frequency (B1) and amplitude (B2) in granule cells one month after FPI. Dotted line illustrates the median. C. Representative current traces illustrate sIPSCs in GCs from rats 3 months after sham (C1) and FPI (C2). D. Cumulative probability plots of granule cell sIPSC frequency (D1) and amplitude (D2) in rats 3 months after FPI. Dotted line illustrates the median.

### Enhancing GABA_A_R δ subunit selective alters GC sIPSC frequency after injury

The decrease in sIPSC frequency 3-5 months after brain injury is consistent with cell-type specific loss of interneurons (Toth et al., 1997; Santhakumar et al., 2000; Frankowski et al., 2019). Since interneuron subtypes differ in expression of GABA_A_R δ subunit mediated tonic GABA currents (Glykys et al., 2007; Yu et al., 2013), we examine whether the proportion of sIPSCs sensitive to THIP were altered 3-5 months after FPI when THIP-induced increase in tonic GABA currents were not different between GCs from controls and FPI rats. In GCs from sham controls, THIP (1 τM) failed to alter sIPSC frequency and amplitude (Fig. 3 A1-D1). Additionally, THIP did not change the 10-90% rise or decay time in GCs in controls indicating limited contribution of GABA_A_R δ subunits to sIPSC kinetics at this time point. In contrast, THIP reduced sIPSC frequency in GCs from rats 3 months after FPI without altering amplitude, τ_decay_ or 10-90% rise time (Fig. 3 A2-D2). The ability of THIP to selectively reduce sIPSC frequency after FPI at a time point when both basal and THIP-induced enhancement of tonic GABA current amplitude were not different between sham and FPI rats suggests that, following brain injury, a greater proportion of synaptic events in GCs are mediated by interneuron subtyprs expressing GABA_A_R δ subunits.

**Figure 3:**
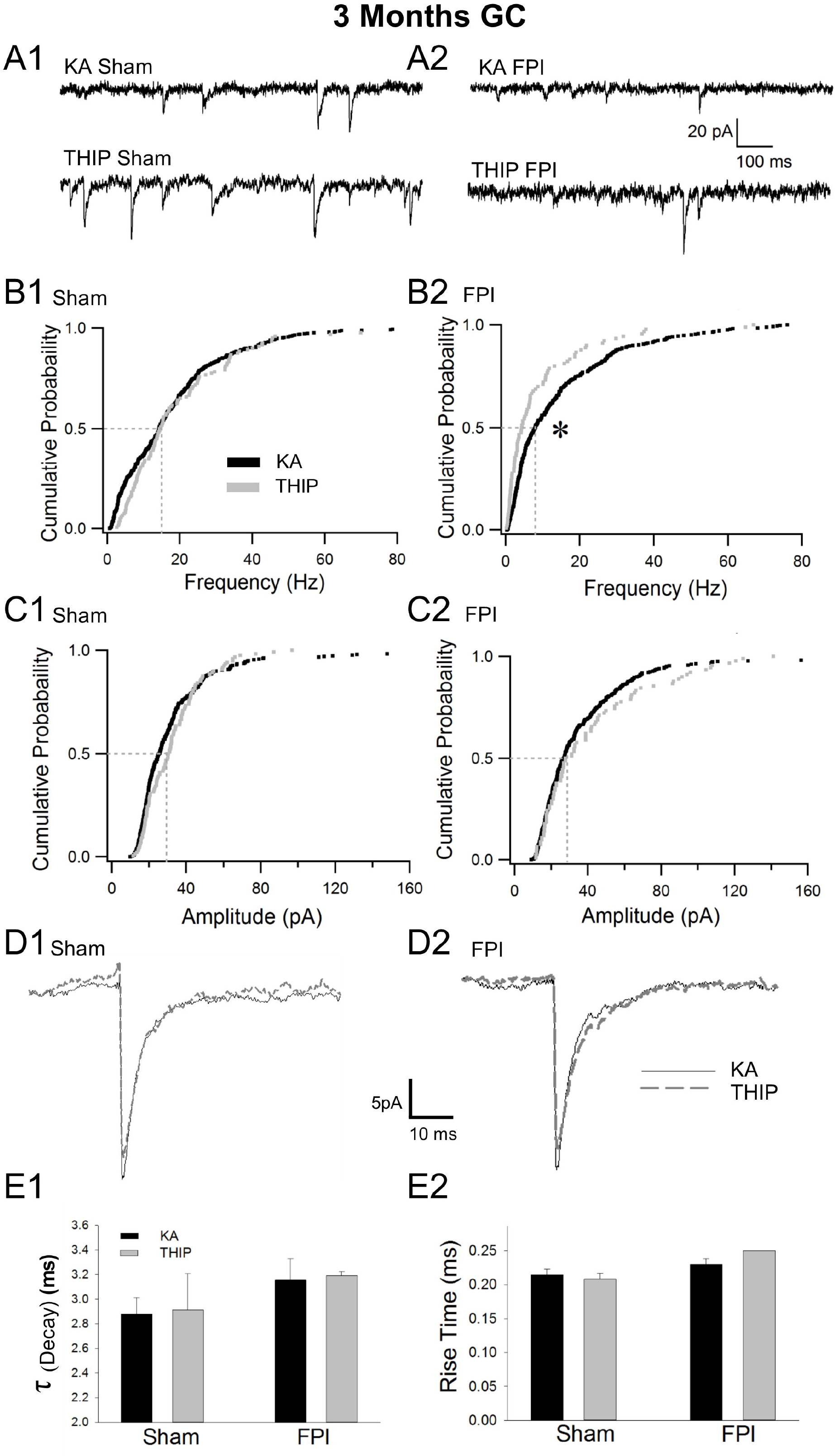
THIP selectively reduces granule cell sIPSC frequency in injured rats. A. Representative current traces illustrate sIPSCs in GCs during baseline recordings in kynurenic acid (above) and in THIP (1 μM, below) in rats three months after sham (A1) and FPI (A2). B. Cumulative probability plots compare sIPSC frequency during baseline recordings and in THIP in sham (B1) and FPI (B2) rats. CB. Cumulative probability plots of sIPSC frequency during baseline recordings and in THIP recorded in sham (C1) and FPI (C2) rats D. Overlay of average sIPSC trace recorded in kynurenic acid and THIP from sham (D1) and FPI (D2) rats. E. Summary histograms of effect of THIP on sIPSC decay time constant (E1) and 10-90 rise time (E2).

### Long term apparent recovery of SGCs inhibition after brain injury

Although SGCs share several structural features with GCs, they show differences in synaptic inputs and sustained firing which make them functionally distinct from GCs (Williams et al., 2007; Larimer and Strowbridge, 2010; Gupta et al., 2020). SGCs undergo a marked decrease in amplitude of tonic GABA currents and sIPSC frequency one-week after FPI, which contrasts with the increase in GCs (Gupta et al., 2012). Since SGCs show developmental reduction in tonic GABA currents (Gupta et al., 2020), we examined whether injury-induced differences in SGC GABA currents persisted to three months. SGCs were identified based on morphological features including multiple primary dendrites, semilunar somata, and wide dendritic span in the molecular later (Fig. 4 A-B). Unlike early after FPI, SGC tonic GABA currents were not different from GCs 3-5 months after FPI (Fig. 4C, D). Similarly, sIPSC frequency and amplitude in SGCs from rats three months after FPI were not different from sham controls (Figure 4F-I; Table 1 and 2). Together, these data demonstrate a sequential progression of changes in synaptic and tonic GABA currents with a decline in inhibition overtime after injury in GCs and an apparent recovery of synaptic inhibition in SGCs.

**Figure 4:**
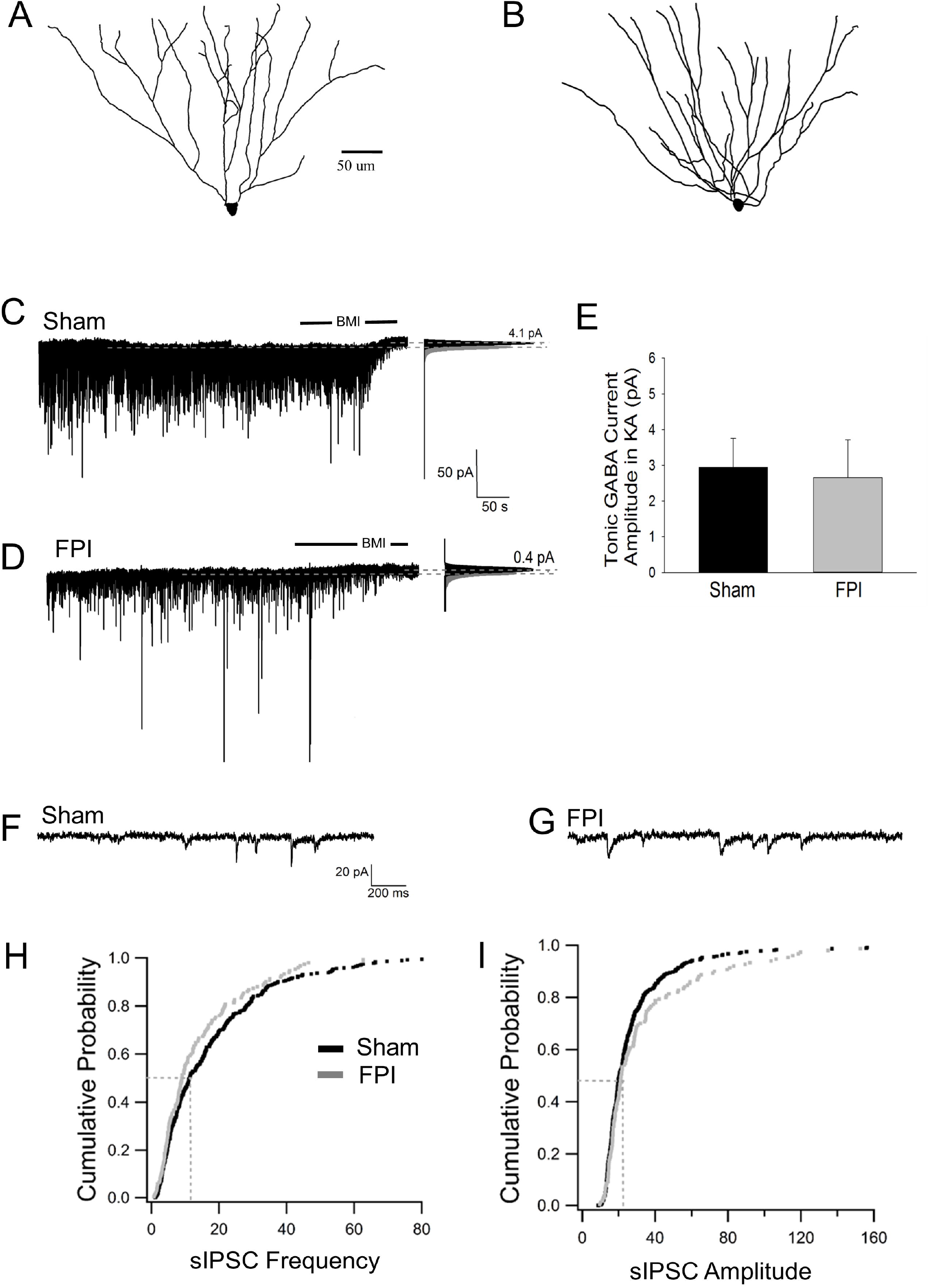
Long-term injury-induced changes in GABA currents in semilunar granule cells. A-B. Representative Neurolucida reconstructions of semilunar granule cells from rats 3 months after sham (A) and FPI (B). C-D. Representative traces from semilunar granule cell three months after sham (C) or FPI (D) illustrate tonic GABA current as the baseline current blocked by bicuculline. Panels to the right show Gaussian fits to the positive half of histograms derived from 30 s recording periods in control conditions and during the perfusion of BMI used to determine tonic current. The dashed lines indicate the Gaussian means, and the difference in currents are noted. E. Summary plot of tonic GABA current amplitude (pA) in SGCs from rats 3 months after FPI. F-G. Representative current traces illustrate sIPSCs in SGCs from rats 3 months after sham (F) and FPI (G). H. Cumulative probability plots of SGC sIPSC frequency (H) and amplitude (I) in rats 3 months after FPI. Dotted line illustrates the median values.

## Discussion

TBI results in diverse cellular and network alteration (Morales et al., 2005; Pitkanen et al., 2009; Hunt et al., 2013; Neuberger et al., 2017a). Acute injury-induced death of GABAergic neurons in the dentate hilus initiates progressive changes in inhibition which differ between injury models (Lowenstein et al., 1992; Toth et al., 1997; Hunt et al., 2011; Pavlov et al., 2011; Frankowski et al., 2019; Parga Becerra et al., 2021). Our data identify progressive changes in tonic and synaptic GABA currents, which differ between the GCs and SGCs. Early post-FPI increase in tonic GABA currents in GCs (Gupta et al., 2012), was absent at three months due to progressive decrease in tonic GABA current in GCs after injury while the amplitude in sham controls remained relatively stable over the same period. In contrast, the apparent recovery of early post-FPI reduction in SGC tonic GABA currents (Gupta et al., 2012), by three months was largely due to a developmental decline in SGC tonic GABA current amplitude in sham rather than a recovery of the injury-induced decrease. Synaptic inhibitory events were examined in the presence of glutamate block to isolate the inhibitory circuit from glutamatergic plasticity (Santhakumar et al., 2001; Hunt et al., 2011; Folweiler et al., 2020). Our results identify that the early posttraumatic increase in sIPSC frequency in GCs (Gupta et al., 2012) gives way to a significant decrease compared to age-matched sham controls by three months. This post-injury decline exceeded the developmental decline in sIPSC frequency observed in GCs (Gupta et al., 2020). Interestingly, THIP selectively reduced GC sIPSC frequency three months post-injury suggesting that a greater proportion of GC sIPSCs in the injured dentate are mediated by inhibitory neurons expressing GABA_A_ receptor δ subunits. In SGCs, sIPSC frequency was not different from age-matched controls three months after injury. Together, these data demonstrate complex, cell-type specific inhibitory plasticity after concussive brain injury in adolescent rats, which evolve alongside developmental plasticity.

Tonic and synaptic GABA currents are known to be altered after brain injury with outcomes varying between experimental models. GC tonic GABA current amplitude was reported to be similar to controls 2-20 weeks after murine CCI or severe concussion in adult rats (Pavlov et al., 2011; Boychuk et al., 2016; Parga Becerra et al., 2021) which contrasts with the increase we had reported one week after moderate concussion in adolescent rats (Gupta et al., 2012). Our current demonstration that GC tonic GABA current amplitude remains elevated one month after injury differs from Pavlov et al. (2011) and parallels our observation at one week (Gupta et al., 2012), while the recover to control levels by three months is consistent with Pavlov et al. (2011). Since, Pavlov et al. (2011) conducted FPI in adult rats, while we adopted FPI in adolescent rats, when tonic GABA current amplitude shows a developmental peak before declining to adult levels (Gupta et al., 2020), age of injury likely contributed to differences in the effect of injury on tonic GABA current amplitude. Differences in injury severity and anesthesia could also contribute to the divergent effects. The lack of change in THIP enhancement of tonic GABA currents after injury indicates that the proportional contribution of δ subunit containing GABA_A_R to tonic GABA currents is retained after FPI and is consistent with Pavlov et al. (2011). It is possible that species or model specific effects underlie the reduction in THIP modulation of GC tonic GABA currents after murine CCI (Boychuk et al., 2016; Parga Becerra et al., 2021). In keeping with interaction between developmental and injury-induced changes, the decrease in SGC tonic GABA currents observed one week post-FPI (Gupta et al., 2012) is eliminated by three months as a consequence of age-related decline in SGC tonic GABA currents in sham controls (reduced from 16.7 ±1.7 pA 1 week after sham injury reported in Gupta et al., 2012, to 2.9 ±0.8 pA) rather than recovery of post-injury changes (4.1±0.9 pA reported 1 week after FPI in Gupta et al., 2012, to 2.7 ±1.1 pA). Thus, brain injury during adolescence may selectively perturb tonic inhibition. Given the differences in posttraumatic pathology between the immature and adult brain (Saletti et al., 2019) and the high incidence of moderate concussions in adolescence, the selective perturbation of tonic GABA currents following concussion during adolescence could contribute to the distinct neuropathology and heightened cognitive deficits in this population (Babikian et al., 2015).

Studies following both murine CCI and severe FPI in rat have reported reduced sIPSC frequency (Pavlov et al., 2011; Boychuk et al., 2016; Parga Becerra et al., 2021), which contrasts with the increase identified in our studies one week after moderate FPI (Santhakumar et al., 2001; Gupta et al., 2012). Since excitatory drive to interneurons can increase after injury (Santhakumar et al., 2001; Hunt et al., 2011; Folweiler et al., 2020), we examined the inhibitory circuit isolated in the presence of glutamate block. Interestingly, the post-injury increase in GC sIPSC frequency observed at one week (Gupta et al., 2012) was no longer evident at one month and reduced further, becoming significantly lower than age-matched controls by three months. In contrast, the decrease in SGC sIPSC frequency observed one week after injury was not present at three months, although the effect reflected developmental decrease in SGC sIPSC frequency in controls (Gupta et al., 2020) rather than recovery of sIPSCs after FPI. Since SGCs are proposed to drive polysynaptic lateral inhibition of GCs (Larimer and Strowbridge, 2010), it is possible that early reduction in SGC inhibition after brain injury may lead to compensatory increase in GC inhibition and impair memory processing, and delay epileptogenesis (Gupta et al., 2012; Folweiler et al., 2020; Korgaonkar et al., 2020b; Korgaonkar et al., 2020a). The maintenance of low SGC inhibition together with decline in GC inhibition could contribute to the enhanced incidence of posttraumatic epilepsy following moderate concussive TBI in adolescent rats (Neuberger et al., 2017b; Korgaonkar et al., 2020b).

Tonic and synaptic inhibition are interrelated, with sIPSCs driving GABA spillover, which augments tonic GABA currents (Glykys and Mody, 2007) while perisynaptically expressed GABA_A_R δ subunits influence IPSC decay kinetics (Wei et al., 2003). Tonic GABA currents also alter baseline noise and conductance, impacting detection of synaptic events and confounding analysis of IPSC frequency and amplitude (Mangan et al., 2005). Parvalbumin-basket cells and molecular layer interneurons express GABA_A_R δ subunits (Glykys et al., 2007; Yu et al., 2013), raising the possibility that enhancing interneuronal tonic GABA currents could reduce interneuron excitability and sIPSC frequency. The lack of injury-induced differences in GC tonic GABA current amplitude and its enhancement by THIP three months after injury enabled us to examine potential impact of THIP on modulation of interneuronal tonic GABA currents on GC sIPSC frequency without the confounding changes in GC conductance or noise. We identified that THIP selectively decreased GC sIPSC frequency after FPI and not in controls, suggesting that THIP may suppress inhibitory neuronal activity after brain injury. While hilar somatostatin neurons undergo extensive cell loss, parvalbumin basket cells and molecular layer interneurons appear to survive (Toth et al., 1997; Santhakumar et al., 2000; Hunt et al., 2011; Frankowski et al., 2019; Folweiler et al., 2020) and may have enhanced contribution to GC sIPSCs after brain injury. It is also possible that interneuronal tonic inhibition is enhanced after brain injury, as observed in epilepsy (Yu et al., 2013), resulting in greater modulation by THIP. Indeed, recent data indicate that inhibition of dentate parvalbumin neurons may be increased after FPI (Folweiler et al., 2020). Together, these data identify a shift in interneuronal mediators of GC synaptic inhibition after FPI with increased contribution from subset of interneurons expressing GABA_A_R δ subunit mediated tonic GABA.

In summary, we find progressive decrease in GC inhibition after concussive brain injury in adolescence while developmental change in SGC inhibition drives the apparent normalization of SGC inhibition. Reduced SGC inhibition could enhance SGC recruitment during afferent inputs impairing its ability to shape input-specific lateral inhibition of GCs degrading memory processing. Progressive decline of the early increases in tonic and synaptic inhibition and eventual depression of sIPSC in GCs three months after FPI could contribute to the alterations in cognitive function and epileptogenesis observed over time after brain injury.

## Acknowledgements

We thank Dr. Luke Fritzky for help with imaging.

## Authorship Confirmation Statement

AG, AP, and FSE performed experiments; AG and FSE analyzed data; AG and VS interpreted results of experiments; AG prepared figures; AG and VS conception and design of research; VS drafted manuscript.

