## Supplemental Table 1 for "Long-term effects of moderate concussive brain injury during adolescence on synaptic and tonic GABA currents in dentate projection neurons"

**TABLE 1: Mean, Median and N**

| | GROUPS | **Tonic GABA Currents** Mean±SE (Median) | **sIPSC Frequency** Mean±SE (Median) | **sIPSC Amplitude** Mean±SE (Median) |
|---|---|---|---|---|
| **Granule Cells (KA)** | | | | |
| 1 Month Post | Sham | 7.2±1.3(6.6) N=8/4 | 17.6±1.1(12.5) N=9/5 | 30.1±1.4(23.0) N=9/5 |
| | FPI | 15.2±2.2(13.9) N=6/5 | 19.4±1.2 (14.9) N=6/5 | 38.5±2.4 (32.3) N=6/5 |
| 3 Months Post | Sham | 5.4±1.4(4.6) N=10/7 | 18.1±0.9 (14.3) N=14/5 | 34.5±1.9 (25.5) N=14/5 |
| | FPI | 7.0±1.8 (6.1) N=9/6 | 14.2±0.8 (8.0) N=10/6 | 40.3±3.6 (27) N=10/6 |
| **Granule Cells (THIP)** | | | | |
| 1 Month Post | Sham | 14.7±3.0 (11.5) N=5/3 | 16.1±1.7 (9.2) N=5/3 | 31.7±1.3 (26.6) N=5/3 |
| | FPI | 32.3±3.4 (32.9) N=5/4 | 25.6±1.8 (21.0) N=5/4 | 19.0±1.3 (14.1) N=5/4 |
| 3 Month Post | Sham | 18.9±3.4 (18.05) N=7/5 | 20.1±1.6 (14.7) N=7/5 | 32.0±1.6 (30.5) N=7/5 |
| | FPI | 20.8±2.8 (22) N=3/3 | 12.7±1.8 (6.6) N=3/3 | 50.4±4.4 (41.8) N=3/3 |
| **Semilunar Granule Cells (KA)** | | | | |
| 3 Month Post | Sham | 2.9±0.8(3.0) N=9/7 | 15.7±1.2(14.4) N=13/6 | 21.1±2.0(19.2) N=13/6 |
| | FPI | 2.7±1.1(2.4) N=6/4 | 13.6±0.8(14.6) N=6/4 | 39.0±9.4(26.6) N=6/4 |
| *N=number of cells/ number of rats* | | | | |
