## Supplemental Table 2 for "Long-term effects of moderate concussive brain injury during adolescence on synaptic and tonic GABA currents in dentate projection neurons"

**TABLE 2: Statistical Analysis**

| GROUPS | **Tonic GABA Currents** (T-Test, p) | **sIPSC Frequency** WRT(p)ED(KST) | **sIPSC Amplitude** WRT(p)ED(KST) | |
|---|---|---|---|---|
| **Granule Cells (Sham vs. FPI)** | | | | |
| 1 Mo KA | *Sig (0.004)* | *Sig ($7.4e^{-9}$)*, No Cohen's D=0.4 | *Sig (0.0003)*, No Cohen's D=0.4 | Figure 1 and 2 |
| 3 Mo KA | NS (0.5) | *Sig ($3.3e^{-6}$)*, No Cohen's D=0.2 | NS (0.33), Yes Cohen's D=0.08 | Figure 1 and 2 |
| 3 Mo THIP | NS (0.75) | Sig ($2.5e^{-13}$), No Cohen's D = 0.7 | *Sig (0.016)*, Yes Cohen's D = 0.2 | Figure 1 and 3 |
| **Granule Cells (KA vs. THIP)** | | | | |
| 3 Mo Sham | *Sig (0.004)* | *Sig(0.007)*, No | NS(0.054), Yes | Figure 3 |
| 3 Mo FPI | NS (0.9) | *Sig(0.0001)*, No | NS (1), No | Figure 3 |
| **Semilunar Granule Cells (Sham vs. FPI)** | | | | |
| 3 Mo KA | NS (0.8) | NS (0.06), Yes Cohen's D = 0.1 | NS (0.08), No Cohen's D = 0.1 | Figure 4 |
| *ED(KST) = Equal Distribution Test (Kolmogorov Smirnov Test); WRT(p) = Wilcoxon Rank Test (p Value)* | | | | |
